# Floral display size does not affect the distribution of paternal diversity in *Silene dioica*

**DOI:** 10.1101/2025.07.19.665651

**Authors:** Nina Joffard, Ihsen Lecat, Kate Ollier, Camille Jolivel, Eric Schmitt, Cécile Godé, Estelle Barbot, Mathilde Dufay, Isabelle De Cauwer

**Affiliations:** Univ. Lille, CNRS, UMR 8198 - Evo-Eco-Paleo, F-59000 Lille, France; Univ. Lille, Plateforme Serres, Cultures et Terrains Expérimentaux, Fed 4129 IRePSE, F-59000 Lille, France; ISEM, Univ. Montpellier, CNRS, IRD, Montpellier, France; CEFE, Univ Montpellier, CNRS, EPHE, IRD, Montpellier, France

**Keywords:** Mating patterns, sire profiles, floral display size, plant-pollinator interactions, paternity analyses, pollen carryover

## Abstract

**Premise of the study:** Paternal diversity and its distribution within maternal plants is governed by various factors, including frequency of pollinator visits, patterns of pollinator movement and pollen carryover. Floral display size may have opposite effects on fruit and plant-level paternal diversity, because plants with large displays should receive more pollinator visits but also more sequential probes per visit.

**Methods:** To investigate the effect of floral display size on the distribution of paternal diversity in *Silene dioica*, we genotyped and assigned paternity for 1320 seeds sampled in several fruits per plant in control and manipulated females, whose flower number was artificially increased. We estimated within-fruit paternal diversity and sire profile dissimilarity among fruits and compared these metrics between control and manipulated plants. We then studied pollen carryover using controlled pollinator visits and pollen counts.

**Key results:** Most fruits were multiply sired, but we found no effect of flower supplementation on within-fruit paternal diversity. Sire profiles were more similar between fruits located on the same than on different females, but sire profile similarity was not higher within manipulated than within control plants. The pollen deposition curve was steep, with an increasing carryover fraction throughout the visitation sequence.

**Conclusions:** Our results suggest that the effect of floral display size on pollinator foraging behaviour is too weak to affect paternal diversity and its distribution in *S. dioica*, or that other processes, such as limited pollen carryover and spatially restricted pollen dispersal, blur the effect of pollinator visit frequency and movements on sire profiles.

## INTRODUCTION

Mating patterns are central to explain the dynamics as well as the evolutionary trajectory of natural populations (Harder and Barrett, 1996; Barrett and Harder, 2017). In flowering plants, mating patterns are shaped by various factors influencing pollen fate, including, in animal-pollinated plants, plant-pollinator interactions (Rhodes et al., 2017). Much of the literature on mating patterns in flowering plants has focused on the balance between self and outcross pollination (Barrett, 2003), while multiple mating has received less attention. Because mating with other individuals requires the intervention of a third agent, the pollen vector, outcrossing in animal-pollinated flowering plants is typically promiscuous. In particular, flowering plants often receive pollen from more than one donor (*i.e.* polyandry). Accordingly, multiple paternity has been documented in various species (reviewed in Bernasconi, 2009; Pannell and Labouche, 2013) and its evolutionary consequences may be significant. The deposition of mixed pollen loads onto individual stigmas – either in the course of a single or during sequential pollinator visits – creates opportunities for pollen competition among donors and female choice, which could both have benefits in terms of offspring quality and maternal fitness (Tonnabel et al., 2021). More generally, multiple paternity should increase offspring genetic diversity, thereby increasing the likelihood of seedling establishment in spatially and temporally heterogenous environments (Karron and Marshall, 1993). Moreover, because plants tend to disperse their seeds locally (Nathan and Muller-Landau, 2000), multiple paternity may affect the genetic diversity of seedlings establishing in the vicinity of maternal plants, and thus, the fine-scale population structure (Wells and Young, 2002).

Because of the modular structure of plants, which often produce multiple flowers, paternal diversity can be distributed in various ways within maternal plants (Pannell and Labouche, 2013). Plants may exhibit comparable levels of overall paternal diversity but differ in the way this diversity is distributed among fruits, *e.g.*, seeds from a given fruit can be sired by multiple pollen donors with overlapping sets of sires across fruits, or conversely, by only one or a few donors with distinct sets of sires across fruits. The distribution of paternal diversity within maternal plant is biologically meaningful, as it can reveal the relative influence of different processes operating at distinct spatial scales. More specifically, within-fruit paternal diversity should depend on the frequency of pollinator visits and on pollen transport dynamics, which respectively influence the likelihood of receiving pollen grains from different donors sequencially and simultaneously (Karron *et al*., 2006; Mitchell *et al*., 2013). Post-pollination processes may further shape this diversity by selectively filtering among pollen grains (Marshall *et al*., 2007; Tonnabel *et al*., 2021). In contrast, among-fruit variation in sire profiles should mainly depend on patterns of pollinator movements and pollen carryover.

Understanding how paternal diversity is distributed within maternal plants is therefore key to identify the mechanisms underlying multiple paternity and its evolutionary implications. Because floral traits partly shape plant-pollinator interactions, they may affect paternal diversity and its distribution within maternal plants. Floral display size, for example, is known to impact both the frequency of pollinator visits and the likelihood of within-plant pollinator movements, and may have contrasting effects on both within and among-fruit paternal diversity. Indeed, plants with large floral displays have repeatedly been shown to be more attractive to pollinators and to receive more visits than plants with few open flowers (Ohashi and Yahara, 2001, 2002; Makino et al., 2007; Nattero et al., 2011). At the same time, optimal foraging theory predicts that pollinators should limit energetic costs by moving between nearby flowers, so that pollinators may sequentially probe more flowers in plants with large floral displays than in plants with few open flowers (Ohashi and Yahara, 2001, 2002; Mitchell *et al*., 2004). In hermaphroditic species, this was shown to increase the fraction of self-pollen deposited onto the stigmas (Harder and Barrett, 1995; Brys and Jacquemyn, 2010; Karron and Mitchell, 2012; Christopher et al., 2021), thereby reducing within-fruit paternal diversity. Although such an effect is excluded in dioecious species, increasing floral display size could make sire profiles more similar among fruits if sequential probes by the same pollinator result in the deposition of similar pollen mixtures (Mitchell *et al*., 2013). However, these expectations may be modulated by the extent of pollen carryover, that is, the fraction of pollen from a given donor that is carried from one recipient flower to the next one. Experimental studies of pollen carryover have shown that the amount of pollen deposited on recipient flowers decreases in a roughly exponential way throughout the visitation sequence, although pollen deposition curves often show a longer tail than predicted by the exponential model, indicating that the carryover fraction is not constant but instead increases throughout the sequence (Morris et al., 1994, 1995). Both the shape of the pollen deposition curve and the rate of pollen decay vary extensively among plant species, depending on various floral characteristics such as flower morphology (Waser and Price, 1984), amount of nectar (Thomson and Plowright, 1980) or type of pollen dispersal units (Harder and Johnson, 2008), but also among pollinator types, depending on their morphology and behaviour (Castellanos et al., 2003; Santa-Martinez et al., 2021). It has been shown that extensive pollen carryover promotes paternal diversity, while in systems where pollen carryover is limited, pollinators tend to transfer pollen from fewer donors on each recipient flower (Mitchell et al., 2013).

Importantly, limited pollen carryover may also mitigate the effect of floral display size on within-fruit paternal diversity, by influencing the extent to which stigmas are saturated with pollen after single visits. Indeed, if stigmas rapidly reach saturation, we do not expect the aforementioned increase in pollinator visit frequency to translate into an increase in the number of sires per fruit. Likewise, limited pollen carryover may mitigate the effect of floral display size on sire profile similarity among fruits. This is because increasing the number of sequential probes per visit should not increase sire profile similarity if pollinators unload most of the pollen they carry on the first few recipient flowers. Quantifying pollen carryover is thus crucial to understand how floral display size affects paternal diversity and its distribution within maternal plants.

While the effect of floral display size on seed set (*i.e.*, female reproductive success) has been well studied (Karron and Mitchell, 2012; Parachnowitsch et al., 2010; Brunet et al., 2021), few studies have explored the relationship between this trait and plant-level paternal diversity (*i.e.*, female mating success). In an experimental population of the dioecious species *Silene dioica*, Barbot et al. (2023) showed that females with large floral displays had a higher mating success (*i.e.*, number of sires, estimated at the plant level) than females with few open flowers. The same trend was observed when comparing females with artificially increased floral displays to control females, suggesting that floral display size affects plant-level paternal diversity through its effect on pollinator attraction (Barbot et al., 2022). However, whether floral display size also affects within-fruit paternal diversity, and more generally, the distribution of such paternal diversity within maternal plants, is unknown.

Here, we build upon the experiment of Barbot et al. (2022) to address this question in *S. dioica*. Using seed genotyping and paternity assignment, we estimated within-fruit paternal diversity (*i.e.*, alpha diversity) as well as sire profile dissimilarity (*i.e.*, beta diversity) among fruits of the same maternal plants to study the effect of floral display size on the distribution of paternal diversity. Our hypothesis was that manipulated plants, which were previously shown to be more attractive to pollinators, would have a higher number of sires per fruit compared to control plants, but because pollinators may visit more flowers on these plants, we predicted that sire profiles would be more similar among fruits from manipulated than from control plants. Because these predictions may be modulated by the extent of pollen carryover, we conducted an additional experiment to build the pollen deposition curve of *S. dioica* when visited by one of its primary pollinators, *Bombus terrestris*.

## MATERIALS AND METHODS

### Study species and experimental material

*Silene dioica* (L.) Clairv. is a perennial herb which is widespread throughout northern and central Europe, where it flowers in late spring-early summer. This dioecious species is sexually dimorphic for flower number and size, with males producing more and larger flowers than females (Kay et al., 1984; Moquet et al., 2020; Barbot et al., 2023). *Silene dioica* is insect-pollinated and has a generalist pollination system with bees and hoverflies being its main pollinators (Barbot et al., 2022).

Plant material was obtained using seeds collected in three forests from northern France. All individuals were sown in a greenhouse and placed in separate 0.7-L pots filled with a standard soil mixture.

### Testing the effect of floral display size on paternal diversity

#### Experimental setup

The experimental setup described below was designed for a previously published study that aimed at estimating selection on flower number and disentangling its different components in *S. dioica* (Barbot et al., 2022). To this end, an array of 172 plants with a 1:1 sex-ratio was kept in an insect-proof greenhouse until the beginning of the experiment, which was conducted in the experimental garden of Lille University, France (50°36’27.9’’N 3°08’36.3’’E), in May-June 2019. No wild population of *S. dioica* was growing in the vicinity. All 172 plants included in the experiment were genotyped for five microsatellites (see Barbot et al., 2022 for details).

At the flowering peak, all 172 plants were placed in a 50 m² experimental plot for 10 days. Plants were randomly assigned to two treatments – control or manipulated – and were arranged according to a grid pattern where sexes and treatments were alternated. Both the control and the manipulated treatments included 43 males and 43 females. In the control treatment, flower number was left unmanipulated, while in the manipulated treatment, floral display size was artificially increased by adding paper-made dummy flowers in order to double the number of flowers displayed by each plant. Several floral traits putatively involved in pollinator attraction (*e.g.*, number of real flowers produced during the experiment, corolla width, calyx height…) were measured over the course of the experiment and pollinator visits were recorded during three observational sessions of 20 minutes for each plant (see Barbot et al. 2022 for details). Female plants used in this experiment produced on average 242.22 (± 67.99) ovules and 157.00 (± 60.18) seeds per flower, and seed set was not pollen limited, as shown by a pollen supplementation experiment. After the experiment, female plants were transferred back to the greenhouse, where fruits were left to mature for a couple of weeks before being collected. Fruits were conserved in individual paper bags at a temperature of 12°C until seed genotyping was performed.

#### Seed genotyping

Among the 86 female plants included in the experiment of Barbot et al. (2022), we selected 10 in the manipulated treatment and 11 in the control treatment. These manipulated and control plants did not differ in terms of floral traits, including total number of real flowers produced during the experiment (F_1,19_ = 0.004, P = 0.948), corolla width (F_1,19_ = 0.03, P = 0.871), calyx height (F_1,19_ = 0.11, P = 0.746) and number of ovules per fruit (F_1,19_ = 0.06, P = 0.814). In each of these plants, we selected two to three fruits and we randomly selected and genotyped 24 seeds per fruit. In total, 1320 seeds from 55 fruits were genotyped. DNA was extracted directly from seeds using the NucleoMag 96 Plant kit (Macherey-Nagel®) (Method S1), and five nuclear microsatellites were amplified and scored as described in Barbot at al. (2022).

#### Paternity assignment

We performed paternity assignment using the maximum-likelihood procedure implemented in CERVUS 3.07 (Marshall et al., 1998; Kalinowski et al., 2007). Likelihood scores, based on allele frequencies measured in the experimental population and on individual genotypes, were calculated for each pair of seed and potential sire, considering as potential sires all 86 males included in the experiment of Barbot et al. (2022) and previously genotyped for the same microsatellites. We used the ΔLOD criterion (i.e. difference in likelihood scores between the two most likely sires) to determine whether seed paternity could be assigned to the most likely sire. The value of ΔLOD below which paternity could not be assigned at 80% was determined using a distribution obtained by simulating 10000 mating events, allowing for a 2% genotyping error.

#### Statistical analyses

All statistical analyses were performed using R version 4.4.2 (R Core Team, 2024). Because the number of seeds whose paternity could be successfully assigned varied among fruits (min = 14, max = 24), we used bootstrapping to sample 14 seeds per fruit 100 times, thereby avoiding biases in the estimation of within-fruit paternal diversity. To assess coverage, we built rarefaction curves for each fruit and calculated the slope of each curve for a sample size of 14 using the *rarify* function from the package *vegan* (Oksanen et al., 2022). To ensure that coverage did not differ between fruits from manipulated and control plants, slopes were then compared between the two treatments using a linear mixed model (LMM) including treatment as fixed effect and maternal plant as a random effect.

The estimated number of sires and effective number of sires per fruit were calculated for each bootstrap and were then averaged across bootstraps. The effective number of sires (Ke) was calculated as 1 / sum(p_i_²), where p_i_ corresponds to the proportion of seeds sired by the i-th pollen donor (Bernasconi, 2009). Note that the effective number of sires thus corresponds to the inverse of the coefficient of correlated paternity, which varies from 0 (all seed pairs are sired by different donors) to 1 (all seed pairs are sired by the same donor) (Ritland, 1989). Both the estimated number of sires and effective number of sires per fruit were then compared between manipulated and control plants using linear models with treatment, number of real flowers and their interaction as fixed effects, and maternal plant as a random effect. The effective number of sires was log-transformed prior to analysis in order to meet the assumption of the linear model. The significance of fixed effects was assessed using Wald chi-square tests.

We then quantified sire profile dissimilarity for each pair of fruits by calculating beta diversity indices following Legendre (2014). We used an incidence matrix with fruits arranged in rows and pollen donors arranged in columns. Dissimilarity coefficients were computed for each pair of fruits, both on a qualitative version of the dataset (with presence/absence of pollen donor *j* in fruit *i*) and on a quantitative one (with number of seeds sired by pollen donor *j* in fruit *i*, *i.e.*, paternity share). For the qualitative analysis, the Sørensen index was computed and decomposed into richness and replacement components following the approach of Podani (Legendre, 2014), to test whether, for each pair of fruits (i) the same number of sires were detected (richness component), and (ii) the same sires were detected (replacement component). For the quantitative analysis, the quantitative version of the Sørensen index (also termed “percentage difference”) was computed. Just as the qualitative one, this index can be decomposed into an “abundance” component, indicating the extent to which fruits differ in terms of total number of seeds, and a “replacement” component, indicating the extent to which they differ in terms of paternity shares. However, because in our case, the number of seeds sampled per fruit was fixed to 14 during bootstrapping, this index was not further decomposed (*i.e.*, because the abundance component would be null for all fruit pairs by definition) (see Method S2 for details). We ran the analysis on each bootstrapped version of the incidence matrix and averaged the resulting dissimilarity coefficients across bootstraps. This resulted in three dissimilarity matrices, containing, for all fruit pairs, the richness component of the Sørensen index, the replacement component of the Sørensen index and the percentage difference.

To assess whether sire profiles were more similar between fruits located on the same than on different females, we performed PERMANOVAs using the function *adonis2* from the package *vegan*, with each of the three dissimilarity matrices as the dependent variable and maternal plant as the independent variable, constraining permutations within treatments. To assess whether sire profiles were more similar among fruits from manipulated than from control plants, we then performed PERMANOVAs within each treatment with maternal plant as the independent variable and calculated F_M-C_obs_, the difference between F values found for the manipulated and control treatments, and compared F_M-C_obs_ to F_M-C_perm_, the difference between F values found for permuted datasets where maternal plants were randomly assigned to one of these two treatments, with 1000 permutations. The p-value was calculated as the proportion of permutations for which F_M-C_perm_ was superior or equal to F_M-C_obs_.

### Quantifying pollen carryover

#### Experimental setup

An experiment was conducted on the campus of the University of Lille in June 2022 to build the pollen deposition curve of *Bombus terrestris*, one of the most frequent pollinator of *S. dioica* (Kay et al., 1984; Goulson and Jerrim, 1997). Bumblebees used in this experiment came from one BioBest® super mini hive containing approximately 40 workers. Bumblebees were trained to forage on *S. dioica* before the experiment and were fed with pollen *ad libitum*. The experimental setup consisted of one male plant and five female plants placed in a flight arena. The male plant was selected to have at least 20 flowers with dehiscent anthers and the female plants between six and ten virgin flowers (individuals were kept in an insect proof greenhouse before the experiment). Each flower on the female plants was marked on the calyx with a different colour using watercolour paint. The female plants were aligned across the width of the tent, numbered from 1 to 5 and covered with a net. A single bumblebee individual was released into the arena and allowed to visit approximately ten male flowers. The male plant was then removed from the arena and the female plants were uncovered to allow for the bumblebee to visit ten female flowers. For each visited flower ordered from the first to the 10th of the visitation sequence, the identity of the female plant and the colour of the paint mark were recorded. Once the visitation sequence was completed, female plants on which at least one flower had been visited were transferred back to the greenhouse to prevent additional visits. This process was repeated with a new bumblebee individual and new sets of male and female plants for subsequent sequences. Each session consisted of five series, and five sessions were conducted in total, each on a different day.

#### Pollen counts

Approximately five hours after the completion of the final series, the stigma of each flower was carefully collected using fine forceps and stored in FAA (ethanol 70%, formaldehyde 35%and acetic acid, 8:1:1). One of the five lobes of each stigma was then rinsed with distilled water, immersed in a bath of sodium hydroxide (NaOH 4 mol.L-1) for 2 hours and rinsed again. Each lobe was then spread across a microscope slide and observed under a fluorescence microscope (Axio Imager 2; ZEISS, Oberkochen, Germany) using white light and a 10x magnification. Panorama images of the stigma lobes were acquired using ZEN software (ZEISS, Oberkochen, Germany) and pollen grains were counted manually on these images. The flower-level pollen load was estimated by multiplying the number of pollen grains detected on one lobe by 5. Pollen loads did not significantly vary among stigma lobes within flowers, as checked on 30 flowers with 3 lobes per flower (repeatability for Poisson-distributed data and 95% CI estimated using the package *rptR* (Stoffel et al., 2017) based on 1000 bootstrap iterations, R = 0.744; 95% CI: 0.552–0.855, p < 0.001).

#### Statistical analyses

After having discarded stigmas on which pollen grains could not be counted and flowers that were visited multiple times during a single series, our final dataset included 186 flowers. To build the pollen deposition curve, we fitted Generalized Linear Mixed Models (GLMMs) to our pollen count data. These models included position of the flower in the visitation sequence as a fixed effect and bumblebee individual as a random effect. Eight models were built, considering either (i) an exponential or a power-law function, (ii) a random intercept or both random intercept and slope for the bumblebee individual effect, and (iii) a Poisson or a negative binomial error structure. Exponential models assume a constant rate of pollen decay throughout the visitation sequence, while power-law models allow for this rate to either increase or decrease through the sequence. A random intercept implies that bumblebee individuals differ in terms of total amount of pollen deposited on female flowers, while a random slope implies that they also differ in terms of rate of pollen decay. Finally, the negative binomial error structures take overdispersion into account compared to the Poisson error structure. We used the function *glmer* from the package *lme4* (Bates and al., 2015) for Poisson models and the function *glmmTMB* from the package *glmmTMB* (Brooke et al., 2017) for negative binomial models. Models were compared based on their AICc and coefficients of the best model were extracted to fit the pollen deposition curve to our pollen count data. One model (power-law model with both a random intercept and a random slope with negative binomial error structure) failed to converge and was thus excluded from the model selection analysis.

## RESULTS

### Within-fruit paternal diversity

Paternity was successfully assigned for a total of 1150 seeds, with a mean of 54.76 (SD = 10.63) seeds per maternal plant.

On the 86 pollen donors included in the experiment, 77 were found to sire at least one seed in the pool sampled here. The siring success of these 77 pollen donors varied from 1 to 75 seeds, with a mean of 14.94 (SD = 14.97) seeds distributed across 1 to 19 fruits.

The estimated number of sires per fruit ranged from 1 to 13, with a mean of 6.29 (± 2.40) sires and most fruits having between 4 and 8 sires (Fig. S1). The effective number of sires per fruit ranged from 1 to 10.53, with a mean of 3.59 (± 1.89) sires and a right-skewed distribution, indicating that paternity was skewed toward few pollen donors (Fig. S1).

Accordingly, in 80% of the fruits, a single pollen donor sired more than one third of the seeds, and its paternity share reached 50% in 23 fruits out of 55. Not all rarefaction curves reached a plateau for N=14 (Fig. S2), but the slope of the curves did not differ between fruits from manipulated and control plants (*W*(1) = 1.66, P = 0.20). There was a positive correlation between the number of flowers reached by any sire and its average paternity share (Pearson correlation coefficient test on log-transformed data: *r*(75) = 0.25, P = 0.03).

We found no effect of treatment, number of real flowers or interaction between the two on the estimated number of sires per fruit, nor on the effective number of sires per fruit (Table 1).

**Table 1.**
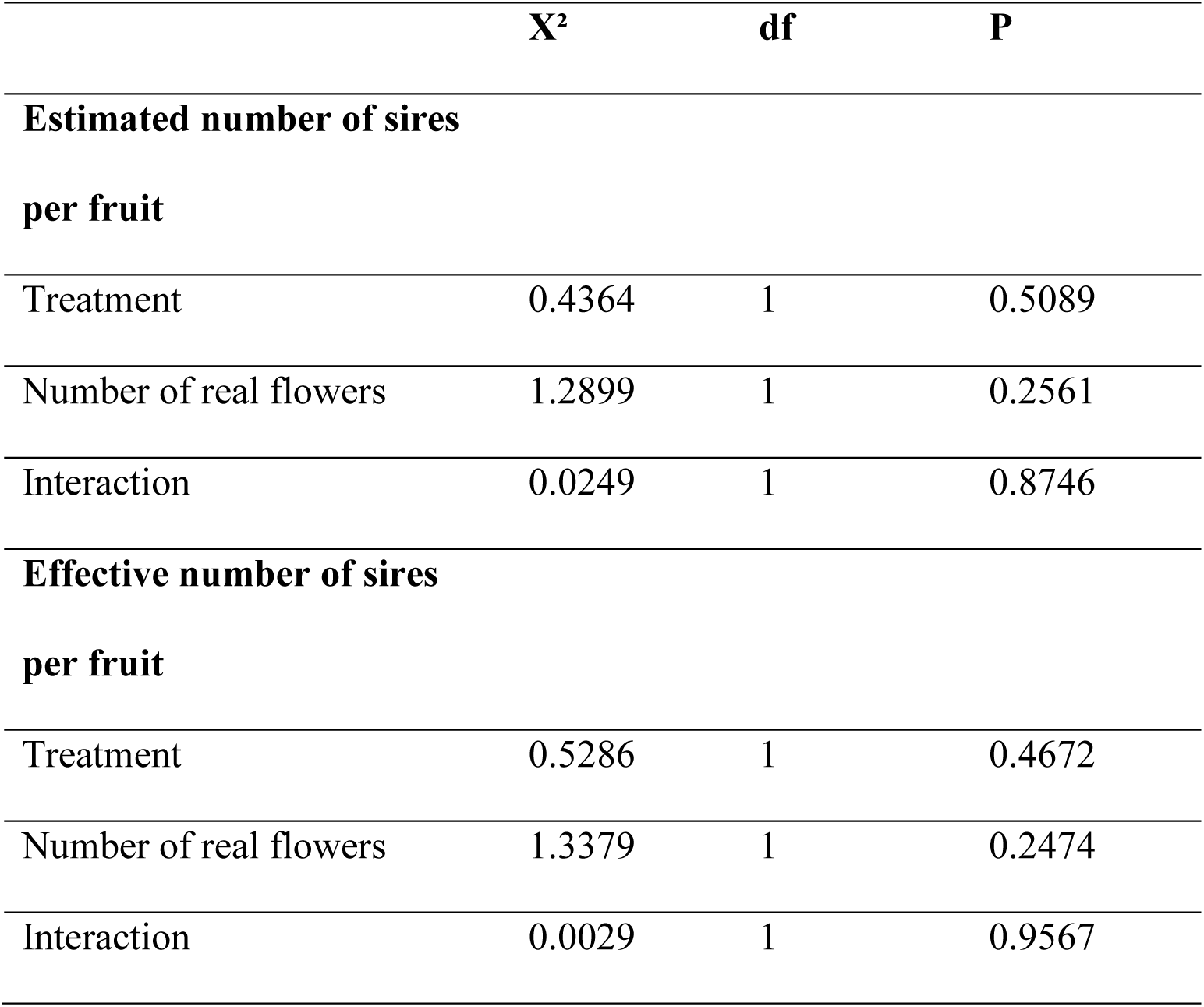
Output of the Wald chi-square tests performed on the coefficients of linear models testing for the effect of treatment, number of real flowers and their interaction on within-fruit paternal diversity.

### Sire profile similarity

The richness component of the Sørensen index ranged from 0.01 to 0.90, with a mean of 0.28. It was significantly higher for fruit pairs located on different maternal plants than for fruit pairs located on the same plant, as shown by the PERMANOVA (F = 2.30, P = 0.005; Fig. 1), indicating that fruits located on the same maternal plant tend to be more similar in terms of paternal diversity than fruits located on different plants. However, this “maternal plant” effect was not stronger in manipulated than in control plants, as indicated by the comparison of F_M-C_obs_ to F_M-C_perm_ (P = 0.55).

**Figure 1.**
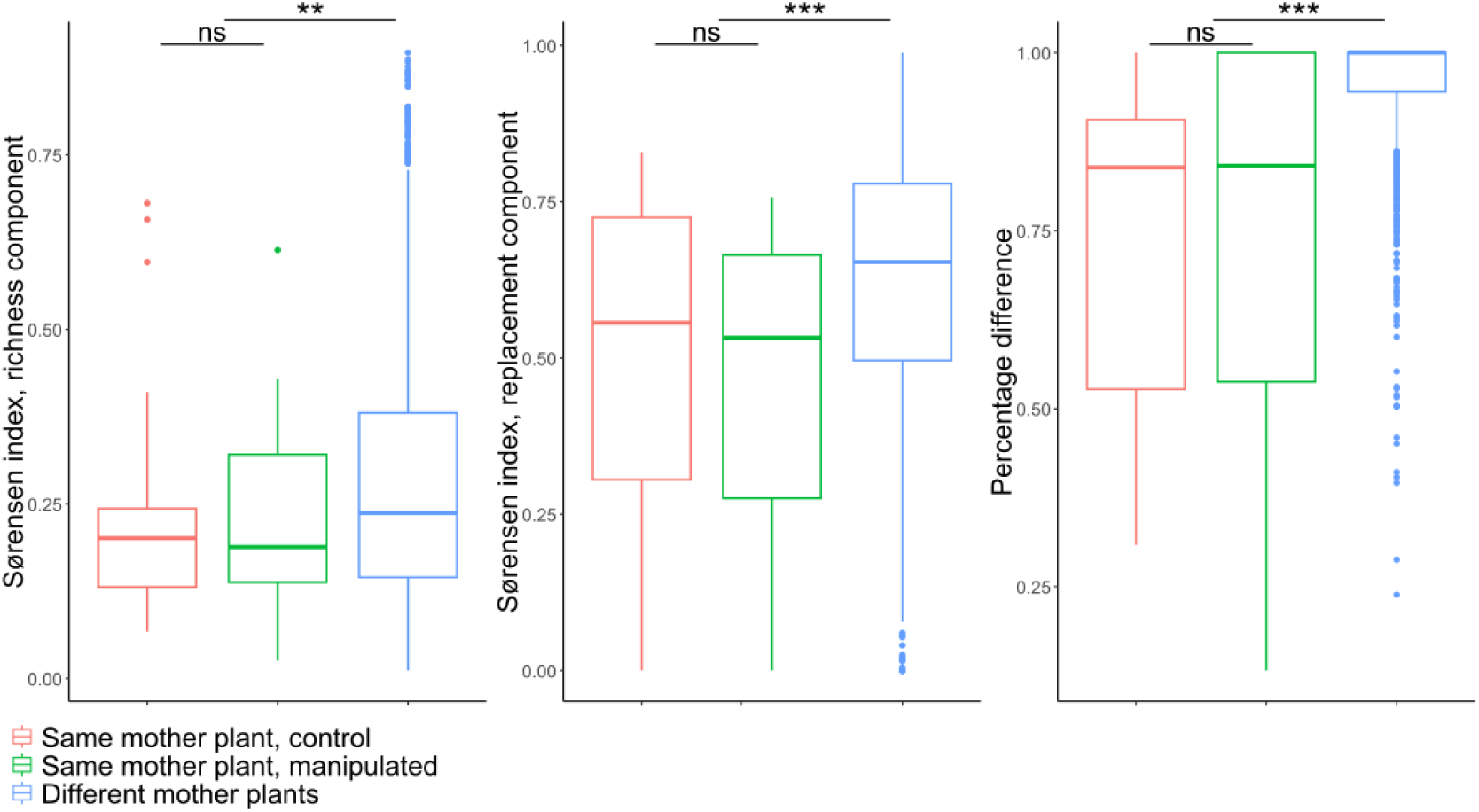
Variation in sire profile dissimilarity between fruit pairs sampled on the same mother plant (control *versus* manipulated) *versus* on different mother plants; significant differences between the two are indicated by asterisks (**: P < 0.01, ***: P < 0.001). The richness component of the Sørensen index indicates variation in the estimated number of sires per fruit, while the replacement component indicates variation in the identity of these sires and the percentage difference variation in their respective paternity shares.

Likewise, the replacement component of the Sørensen index was significantly higher for fruits located on different maternal plants than for fruits located on the same plant (F = 2.36, P = 0.001; Fig. 1), indicating that we detected the same sires more often in fruits sampled on the same maternal plant than in fruits sampled on different plants. Accordingly, fruits located on the same maternal plant shared on average 1.9 sires, against 0.7 for fruits located on different plants. Nevertheless, fruits from manipulated plants did not show more similar qualitative sire profiles than fruits from control plants (P = 0.10).

The same pattern was observed when looking at paternity shares. The percentage difference had a mean of 0.73 for fruits located on the same maternal plant, against 0.95 for fruits located on different plants, and the PERMANOVA indicated that fruits sampled on the same maternal plant shared the same major sires more often than fruits sampled on different plants (F = 2.37, P = 0.001; Fig. 1). Accordingly, in almost half of the maternal plants, the major sire was the same in at least two fruits out of three. However, fruits from manipulated plants did not show more similar quantitative sire profiles than fruits from control plants (P = 0.41), indicating no effect of flower supplementation on sire profile similarity.

### Pollen deposition curve

Negative binomial models performed better than Poisson models, due to overdispersion in our pollen count data. Among negative binomial models, the power-law model performed better than the exponential ones, and models considering a random intercept performed better than those considering both a random intercept and a random slope (Table 2). Thus, the best model (AICc = 1930.2) was a power-law model considering a random intercept only with a negative binomial error structure. According to this model, pollinators deposited on average 398 pollen grains on the first flower of the visitation sequence, and 398 * x^-1.223^ pollen grains on subsequent flowers, where x represents the order of the flower in the visitation sequence (Fig. 2). In other words, according to this model, 41% of the total number of pollen grains delivered across the 10-flowers sequence were deposited on the first flower, and over three quarters were delivered to the first four flowers, while after the seventh flower, less than 10% remained to be deposited.

**Figure 2.**
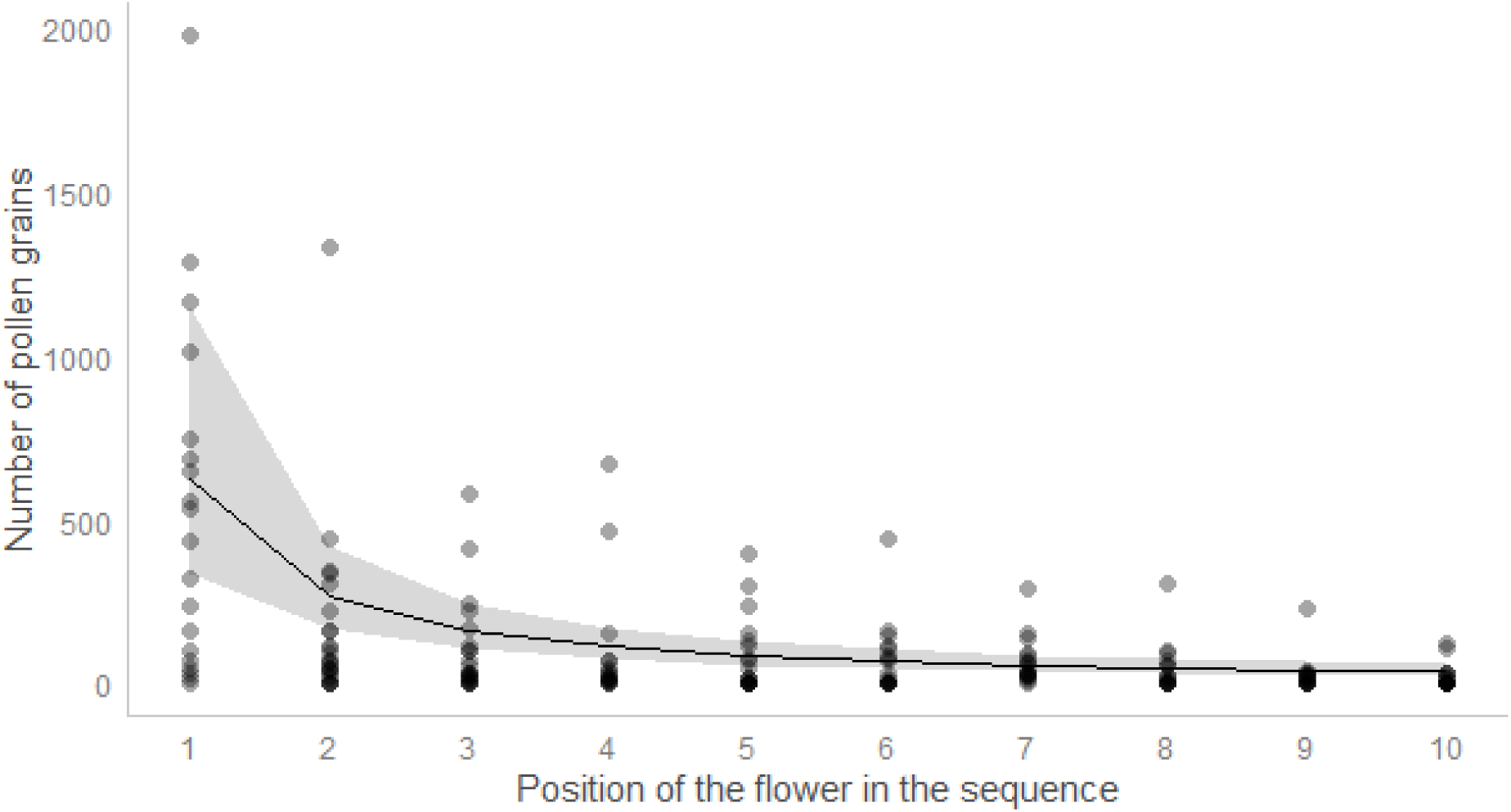
Pollen deposition curve of *S. dioica* when visited by *B. terrestris*. The solid line corresponds to the best model, *i.e.* power-law model with random intercept and negative binomial error structure.

**Table 2.** Description of the models fitted to the pollen count data and corresponding, ranked by increasing AICc.

| **Pollen decay function** | **Bumblebee**<br>**individual effect** | **Error**<br>**structure** | **AICc** |
| --- | --- | --- | --- |
| Power law | Random intercept | Negative<br>binomial | 1930.2 |
| Exponential | Random intercept | Negative<br>binomial | 1934.5 |
| Exponential | Random intercept<br>+ slope | Negative<br>binomial | 1937.9 |
| Power law | Random intercept<br>+ slope | Poisson | 15081.4 |
| Exponential | Random intercept<br>+ slope | Poisson | 15820.0 |
| Power law | Random intercept | Poisson | 19734.3 |
| Exponential | Random intercept | Poisson | 22182.8 |

## DISCUSSION

Paternal diversity and its distribution within maternal plants depend on various factors, including pollinator foraging behaviour and pollen transport dynamics. In this study, we focused on the effect of floral display size of recipient plants, expecting a higher number of sires per fruit, but also a higher sire profile similarity among fruits in recipient plants with large displays, which were previously shown to be more attractive to pollinators (Barbot *et al*., 2022). Contrary to these expectations, we found no effect of flower supplementation, neither on within-fruit paternal diversity nor on sire profile similarity between fruits. Hereafter, we discuss possible explanations for these results.

### Multiple paternity is common in *S. dioica*

In line with other studies (Pannell and Labouche, 2013), our results show that multiple paternity was common in our experimental population, as almost all fruits were multiply sired with an average of 6.29 (SD = 2.40) sires per fruit, for an average of 242 ovules. In the closely related *Silene latifolia*, the estimated number of sires per fruit ranged from 1 to 9, with population means ranging from 3.4 to 4.9 (Teixeira and Bernasconi, 2007). Comparable values were found in species with either much less or much more ovules per flower (e.g., *Ipomopsis aggregata*, a dozen; *Mimulus ringens*, several thousand), but note that variation among studies in the number of genotyped offspring is limited, with this number rarely exceeding 20 per fruit (Pannell and Labouche, 2013), which imposes an upper limit on the number of sires that can be detected. Here, we initially sampled 24 and randomly subsampled 14 seeds per fruit, which represents less than 10% of the total number of seeds produced.

Because in most fruits paternity was highly skewed toward a few pollen donors, increasing our sample size may have only marginally changed our results. In contrast, in some cases paternity shares were more evenly distributed among pollen donors, so that increasing our sample size may have allowed to detect more sires. Importantly, coverage did not differ between manipulated and control plants, enabling a robust comparison of within-fruit paternal diversity between the two treatments.

One important question regarding multiple paternity is whether the latter arises from simultaneous deposition of mixed pollen loads or from sequential deposition of pollen from different donors. Bumblebee foraging behaviour and limited carryover in *M. ringens* suggest that multiple paternity in this species is due to sequential deposition, each pollinator delivering pollen from 1 to 3 donors only (Mitchell et al., 2005; Karron et al., 2006). The opposite conclusion was reached in *I. aggregata*, with multiple paternity being primarily attributed to the extensive pollen carryover observed in this hummingbird-pollinated species (Campbell, 1998). In *S. dioica*, the relative importance of these two processes depends on the extent to which the stigma is saturated after the first visit, which in turn depends on the order of the focal flower in the visitation sequence. Indeed, our pollen carryover experiment showed that when a female flower is visited after several male flowers, it will typically receive hundreds of pollen grains, which should be enough to fertilize all of its ovules, or at least to cause the wilting of the stigma, as shown in *S. latifolia* (Burkhardt et al., 2009). In this case, all paternal diversity should stem from a single pollination event (*i.e.*, simultaneous deposition). However, our experiment also showed that the pollen deposition curve of bumblebee-pollinated *S. dioica* plants was steep, suggesting that when a female flower is visited after several female flowers, it will likely receive few pollen grains, creating an opportunity for subsequent floral visitors to deposit additional pollen grains.

### Within-fruit paternal diversity does not increase with floral display size

We found no differences in the number of sires per fruit between manipulated and control plants, despite the fact that manipulated plants received more pollinator visits and had a higher mating success at the plant level (Barbot et al., 2022). Likewise, in *M. ringens*, the number of sires per fruit did not increase with floral display size (Mitchell et al., 2005), even though plants with large displays were more attractive to pollinators (Mitchell et al., 2004). In this case, this could be explained by the fact that pollinators also visited more flowers per plant in plants with large displays, thereby increasing the fraction of self-pollen deposited onto the stigmas (Mitchell et al., 2005). In our case, geitonogamous pollination cannot explain why the number of sires per fruit does not increase with the number of open flowers, but hereafter we discuss several alternative explanations for this finding.

First, in cases where multiple paternity is primarily due to simultaneous deposition, within-fruit paternal diversity should depend more strongly on the sequence (i.e., order of visits on male and female flowers) than on the number of pollinators visits. In the same species, Moquet et al. (2022) suggested that the number of pollen grains deposited onto a given stigma depends more strongly on the number of male flowers visited before the female flower is reached than on the number of pollinator visits that the latter receives. Likewise, we suspect that within-fruit paternal diversity critically depends on the number of male plants visited before a given female flower is reached. Moreover, in cases where multiple paternity is primarily due to sequential deposition, more pollinator visits may not necessarily imply more pollen donors if females repeatedly receive pollen from the same subset of males, as expected when pollen transfer networks are spatially structured. In line with this hypothesis, Barbot et al. (2022) documented spatially restricted pollen dispersal in this experimental population, showing that the average pollen dispersal distance was 1.50 m (SD = 0.32). Alternatively, more pollinator visits may imply more pollen donors but not necessarily more sires if the first pollen grains to reach the stigma preempt most ovules, as observed in *S. latifolia* (Burkhardt et al., 2009), or if variation in terms of germination or pollen tube growth rates among donors leads to nonrandom patterns of ovule fertilization (Marshall et al., 2007; McCallum and Chang, 2016).

#### Paternity is correlated at the plant level, but not more so in manipulated than in control plants

Our results indicate that sire profiles are more similar between fruits collected on the same maternal plants than between fruits collected on different maternal plants. More specifically, we both found the same sires, and the same major sires more often in fruits from the same plant than in fruits from different plants, indicating correlated paternity at the plant level.

Likewise, correlated paternity was high both within and among fruits in *Eichhornia paniculata* (Morgan and Barrett, 1990), while in *Eucalyptus rameliana*, correlated paternity was high within, but not between fruits, reflecting limited mate availability at any one time but high turnover in available mates during the flowering season (Sampson, 1998). Correlated paternity at the plant level can either result from within-plant pollinator movement, where the same insect visits several flowers on the same plant and hence, deposits the same pollen mixture, or from spatially restricted pollen dispersal, where females tend to repeatedly receive pollen from their male neighbors. A reanalysis of the data collected by Barbot et al. (2022) indicates that during the experiment, pollinators probed on average 1.3 (SD = 0.5) flowers per plant in females, against 2.9 (SD = 2.0) in males. This, combined with the limited pollen carryover evidenced in this study, suggests that in our system, correlated paternity at the plant level primarily reflects the spatially restricted pollen dispersal previously demonstrated in this system (Barbot et al., 2022). Furthermore, we did not find an increased similarity in sire profiles between fruits from the same maternal plants in manipulated compared to control plants. This suggests that the number of flowers probed per plant does not differ between manipulated and control plants, or at least that this difference is too small to cause a detectable difference in sire profile similarity. Accordingly, a reanalysis of the data collected by Barbot et al. (2022) indicates that during the experiment, pollinators probed on average 1.30 (SD = 0.60) flowers per plant in manipulated females, against 1.27 (SD = 0.42) in control ones. We thus hypothesize that adding dummy flowers to *S. dioica* plants increased their detectability and attractivity towards pollinators without affecting patterns of within and among-plant movements.

### Pollen carryover is limited in bumblebee-pollinated *S. dioica* plants

Studying pollen transport dynamics is crucial to understand how patterns of pollinator movements translate into patterns of pollen – and ultimately gene – flow. The pollen deposition curve built in this study was steep, as previously shown in other bee-pollinated species (Castellanos et al., 2003; Santa-Martinez et al., 2021). Together with pollinator foraging behaviour, this limited pollen carryover likely shapes mating patterns in *S. dioica* populations. In particular, it likely contributes to the spatially restricted pollen dispersal and fine-scale population structure reported in previous studies (Ingvarsson and Giles, 1999; Barbot et al., 2022). The pollen deposition curve showed a longer-than-exponential tail due to an increasing carryover fraction throughout the visitation sequence, as found in other systems (Morris et al., 1994). Several mechanisms could explain this, including the early deposition of weakly adhering pollen grains, whereas more strongly adhering grains tend to remain on the pollinator until late in the sequence (Morris et al., 1994). This may result in a pattern where males export most of their pollen to neighboring females while still delivering small amounts of pollen to more distant ones, potentially acting as major sires in the former, and as minor sires in the latter. Our results further show that adding a random “bumblebee individual” intercept, but not slope improved model fit, suggesting that bumblebee individuals varied in the total amount of pollen they carried, but not in the rate at which pollen was depleted from their body. Note that there was still overdispersion in our data afterwards, indicating variation among flowers within a single sequence, which could be due to variation in the duration of pollinator visits - which we did not control for in this study - or to other factors affecting the efficiency of pollen transfer (Richards et al., 2008). This highlights the need for additional experiments to gain a more complete understanding of pollen transport dynamics and resulting mating patterns in *S. dioica*. In particular, although bumblebees are major pollinators of *S. dioica*, they are not the only ones; thus, quantifying pollen carryover in other insects, such as hoverflies, would be valuable to understand how different pollinators contribute to pollen dispersal in this species. Moreover, a comprehensive understanding of pollen transport dynamics would require studying the dynamics of pollen removal, and not only that of pollen deposition. In particular, it would be interesting to assess whether mechanisms such as pollen layering might bias paternity towards the most recently visited males (Minaar *et al*., 2019; Moir & Anderson, 2023).

## CONCLUSIONS

In this study, we showed that although most fruits were multiply sired in our experimental population of *S. dioica*, with an average of 6 sires per fruit, paternity was skewed toward a few pollen donors, which sired a large proportion of the seeds within a fruit. Paternity was also correlated at the plant level, as sire profiles were more similar between fruits located on the same than on different plants. Such patterns of correlated paternity may arise from pollen codispersion, where pollen originating from the same males is deposited on the same flower and on flowers of the same plant during a single pollinator visit, but also from spatially restricted pollen dispersal, with females repeatedly receiving pollen from their male neighbors. We found no effect of flower supplementation on within-fruit paternal diversity even though manipulated plants were previously shown to be more attractive to pollinators, perhaps because the number of sires per fruit depends more strongly on the sequence than on the number of pollinator visits. Likewise, we found no effect of flower supplementation on sire profile similarity among fruits, suggesting that floral display size has little effect on the number of probes per visit. The pollen deposition curve of bumblebee-pollinated *S. dioica* plants was steep, in line with the spatially restricted pollen dispersal previously documented in this species. Further studies are needed to understand how pollinator foraging behaviour, pollen transport dynamics and post-pollination processes interact to shape mating patterns in *S. dioica* populations, ultimately influencing gene flow and population structure.

## ACKNOWLEDGEMENTS

This work has been performed using infrastructure and technical support of the "Plateforme Serre, cultures et terrains experimentaux - Universite de Lille" for the greenhouse/field facilities. We are grateful to Laurence Debacker and Sylvie Flourez for laboratory assistance, and to A. Duputié and N. Hautekeete for helpful discussions. The study was financially supported by the French Agence Nationale de la Recherche through the ANR-JCJC-EvoPoD (ANR-21-CE02-0011).

## AUTHOR CONTRIBUTIONS

NJ and IDC conceived the study questions; NJ, IL, KO, CJ, ES, CG, EB, MD and IDC collected the data; NJ, IL, KO and IDC analyzed the data; NJ led the writing of the manuscript. All authors contributed to revisions and gave final approval for publication.

## DATA AVAILABILITY STATEMENT

Data will be made available on Dryad (http://datadryad.org/).

