## Supplementary Material for "Floral display size does not affect the distribution of paternal diversity in *Silene dioica*"

**Method S1. Seed genotyping in *Silene dioica*.**

To perform paternity analysis in flowering plants, it is standard practice to genotype seedlings, which requires germinating the seeds before DNA exactions and PCR amplifications **(e.g.Meagher, 1986, Broyles and Wyatt, 1990).** This step is considered as necessary, because one may not be able to obtain the offspring's genotype by genotyping seeds directly, due to their structure. Indeed, seeds are formed through double fertilization, where two sperm cells fertilize the haploid egg and the binucleate central cell of the female gametophyte. Seeds thus typically consist of tissues with three distinct genomic compositions:

1. the fusion of the first haploid sperm cell and the haploid egg forms a diploid zygote which will become the **embryo** (2N)
2. the fusion of the second haploid sperm cell (which originates from a mitosis of the same haploid cell as the first and shares the same genetic content) and the two polar nuclei of the central cell of the female gametophyte (which arise by mitosis from the same single meiotic product that gave rise to the egg) forms a triploid primary endosperm cell which will become the **endosperm** (3N)
3. the outer protective layer, or **seed coat**, which develops from the integuments of the ovule and is therefore composed of maternal sporophytic tissue (2N)

*Silene* species have “type P” seeds, where the embryo is peripheral, adjacent to the seed coat, and typically large and elongated (Martin, 1946, Baskin and Baskin, 2019). The embryo is wrapped around a well-developed **perisperm** (a nutritive tissue derived from the nucellus, *i.e.* maternal sporophytic tissue, 2N), while the **endosperm is reduced in mature seeds (Mohana Rao et al., 1988 , Kartal and Tekin, 2021).**

Genotyping whole seeds in *Silene* may thus pose a risk of being unable to isolate the offspring’s genotype due to the mixing of the embryonic tissue (and the endosperm, which reflects the same genetic contribution) with tissues of maternal-sporophyte origin (seed coat and perisperm). This risk depends not only on the relative size of the embryo compared to the other tissues within the seed, but also on the potential degradation of maternal tissues in mature seeds. In particular, DNA contained in maternal tissues such as the seed coat and perisperm may degrade over time. To check whether it is possible to determine the genotype of an offspring by genotyping the seed directly instead of the seedling, we performed a series of controlled crosses using hand pollinations where the dehiscent anthers of one male were rubbed against the receptive stigma of one flower on one female and genotyped a sample of the seeds resulting from these crosses using microsatellites that were previously successfully used to perform paternity analysis in *Silene dioica* (see Barbot et al., 2022, Barbot et al., 2023, Barbot et al., 2025). A total of 12 couples, as well as eight seeds per couple, were genotyped (N = 96 seeds). We then examined the obtained seed genotypes with special attention to (i) parent / offspring genotype discrepancies, (ii) occurrence of triploid genotypes, which would be indicative of maternal tissue contamination and (iii) relative frequencies of homozygotes and heterozygotes in offsprings compared to what would be expected under simple Mendelian segregation given the parents’ genotypes.

***DNA extraction and microsatellite genotyping -*** Total genomic DNA of adult plants was extracted from dried leaf tissue and then subjected to PCR amplification of five nuclear microsatellite markers following the protocol described in Barbot *et al*. (2022). The seeds were ground to a fine powder in individual tubes containing three steal beads each (MN beads, type D, Macherey-Nagel®) using a laboratory tissue grinder (MM400, Retsch®) for 1 min at 30 Hz. Total genomic DNA of seeds was then extracted using the Nucleomag plant Kit (Macherey-Nagel) and an automated DNA purification workstation (KingFisher Flex, Thermo Fisher Scientific, Waltham, USA). PCR amplification and genotyping were performed as described for adult leaves (Barbot et al., 2022).

***Results*** - DNA from all but two seeds was successfully extracted (with a concentration ranging from 1 to 2 ng/µL) and genotyped (94 genotypes, see Table S1). Only one parent / offspring discrepancy was detected (couple 4, seed 7, locus Sd05, see Table S1). We detected no triploid genotypes in seeds, which would have been indicative of a mix of maternal sporophytic tissue with the embryo tissue. The probability of detecting triploids in this scenario obviously depends on the level of polymorphism and on the alleles carried by both parents. If maternal contamination occurs, it would necessarily be detected if both parents are heterozygous for distinct sets of alleles or if the mother is heterozygous and the father homozygous for an allele not carried by the mother. These configurations were found multiple times in our dataset and never resulted in triploids (see shaded cells in Table S1). This result is further supported by the data presented in the main body of the article: out of the 1320 genotyped seeds, none exhibited triploidy, despite the high polymorphism observed in the experimental population (91 alleles in total, with 66, 11, 3, 3, and 8 alleles for Sd05, Sd19, Sd23, Sd25, and Sn03, respectively). Finally, we compared the number of heterozygous and homozygous offsprings observed in our dataset to that expected given the genotypes of both parents in our controlled crosses using a chi-square goodness-of-fit test. If maternal contaminations occurred, even in situations that would not necessarily create visible triploidy (non-shaded cells in Table S1), they should still inflate the number of heterozygotes. This does not appear to be the case here, as our data do not support any differences between the observed and expected numbers for any of the five loci used in our study (see Table S1).

***Conclusions -*** Taken together, these results strongly support that DNA extraction and microsatellite genotyping of whole seeds in Silene dioica allows for the isolation of the offspring’s genotype without being affected by potential contamination from maternal sporophytic tissues (seed coat and perisperm). This may be explained by DNA degradation in these maternal tissues in mature seeds, which should be further investigated by comparing genotyping results from seeds at different maturity stages. Our results thus support that paternity analyses in Silene dioica can be conducted without the need to germinate seeds to obtain seedlings.

**Table S1** - Multilocus genotypes for adults (12 males and 12 females) and for eight seeds per couple. The star indicates the only parent / offspring discrepancy. Shaded cells show situations where a maternal contamination would result in the production of an apparent triploid genotype when scoring seeds (*i.e.* when both parents are heterozygous for distinct sets of alleles or when the mother is heterozygous and the father is homozygous for an allele not carried by the mother). No triploids were observed. The last two rows of the table show the results of the chi-square goodness-of-fit test, assessing whether the observed and expected numbers of heterozygous and homozygous offspring differ.

|  | **Marker** | **Sd05** |  | **Sd19** |  | **Sd23** |  | **Sd25** |  | **Sn03** |  |
| --- | --- | --- | --- | --- | --- | --- | --- | --- | --- | --- | --- |
| **Couple 1** | **Female 1** | **172** | **183** | **143** | **156** | **221** | **224** | **173** | **173** | **216** | **218** |
|  | **Male 1** | **219** | **230** | **143** | **151** | **224** | **224** | **174** | **176** | **218** | **218** |
|  | **Seed 1** | 183 | 219 | 151 | 156 | 224 | 224 | 173 | 176 | 216 | 218 |
|  | **Seed 2** | 172 | 230 | 143 | 151 | 224 | 224 | 173 | 174 | 218 | 218 |
|  | **Seed 3** | 183 | 219 | 143 | 151 | 224 | 224 | 173 | 176 | 216 | 218 |
|  | **Seed 4** | 172 | 219 | 151 | 156 | 224 | 224 | 173 | 176 | 216 | 218 |
|  | **Seed 5** | 172 | 219 | 143 | 143 | 224 | 224 | 173 | 176 | 216 | 218 |
|  | **Seed 6** | 183 | 230 | 143 | 151 | 224 | 224 | 173 | 174 | 218 | 218 |
|  | **Seed 7** | 172 | 219 | 143 | 156 | 224 | 224 | 173 | 176 | 218 | 218 |
|  | **Seed 8** | 172 | 219 | 143 | 143 | 224 | 224 | 173 | 176 | 218 | 218 |
| **Couple 2** | **Female 2** | **178** | **197** | **143** | **149** | **221** | **221** | **173** | **173** | **216** | **218** |
|  | **Male 2** | **197** | **201** | **143** | **143** | **221** | **224** | **173** | **173** | **216** | **218** |
|  | **Seed 1** | 197 | 201 | 143 | 149 | 221 | 224 | 173 | 173 | 216 | 216 |
|  | **Seed 2** | 178 | 201 | 143 | 143 | 221 | 224 | 173 | 173 | 216 | 218 |
|  | **Seed 3** | 178 | 197 | 143 | 149 | 221 | 221 | 173 | 173 | 216 | 218 |
|  | **Seed 4** | 197 | 201 | 143 | 143 | 221 | 224 | 173 | 173 | 216 | 216 |
|  | **Seed 5** | 178 | 201 | 143 | 143 | 221 | 221 | 173 | 173 | 216 | 216 |
|  | **Seed 6** | 178 | 201 | 143 | 143 | 221 | 224 | 173 | 173 | 216 | 218 |
|  | **Seed 7** | 178 | 201 | 143 | 143 | 221 | 224 | 173 | 173 | 216 | 216 |
|  | **Seed 8** | 197 | 197 | 143 | 143 | 221 | 221 | 173 | 173 | 218 | 218 |
| **Couple 3** | **Female 3** | **178** | **188** | **143** | **151** | **221** | **224** | **173** | **174** | **218** | **218** |
|  | **Male 3** | **165** | **174** | **143** | **143** | **224** | **224** | **173** | **173** | **216** | **218** |
|  | **Seed 1** | 165 | 178 | 143 | 143 | 221 | 224 | 173 | 174 | 218 | 218 |
|  | **Seed 2** | 165 | 188 | 143 | 151 | 224 | 224 | 173 | 174 | 218 | 218 |
|  | **Seed 3** | 174 | 178 | 143 | 143 | 224 | 224 | 173 | 174 | 218 | 218 |
|  | **Seed 4** | 174 | 188 | 143 | 151 | 224 | 224 | 173 | 174 | 216 | 218 |
|  | **Seed 5** | 174 | 188 | 143 | 151 | 224 | 224 | 173 | 174 | 216 | 218 |
|  | **Seed 6** | 174 | 178 | 143 | 151 | 221 | 224 | 173 | 174 | 216 | 218 |
|  | **Seed 7** | 165 | 178 | 143 | 151 | 221 | 224 | 173 | 174 | 218 | 218 |
|  | **Seed 8** | 165 | 178 | 143 | 143 | 221 | 224 | 173 | 174 | 216 | 218 |
| **Couple 4** | **Female 4** | **166** | **166** | **143** | **143** | **224** | **227** | **173** | **173** | **220** | **220** |
|  | **Male 4** | **186** | **198** | **143** | **143** | **221** | **224** | **173** | **173** | **218** | **220** |
|  | **Seed 1** | 166 | 186 | 143 | 143 | 224 | 224 | 173 | 173 | 218 | 220 |
|  | **Seed 2** | 166 | 198 | 143 | 143 | 221 | 224 | 173 | 173 | 220 | 220 |
|  | **Seed 3** | 166 | 198 | 143 | 143 | 224 | 224 | 173 | 173 | 220 | 220 |
|  | **Seed 4** | 166 | 198 | 143 | 143 | 224 | 227 | 173 | 173 | 220 | 220 |
|  | **Seed 5** | 166 | 186 | 143 | 143 | 224 | 227 | 173 | 173 | 220 | 220 |
|  | **Seed 6** | 166 | 198 | 143 | 143 | 221 | 227 | 173 | 173 | 218 | 220 |
|  | **Seed 7** | 166 | 200* | 143 | 143 | 224 | 227 | 173 | 173 | 220 | 220 |
|  | **Seed 8** | 166 | 198 | 143 | 143 | 224 | 227 | 173 | 173 | 220 | 220 |
| **Couple 5** | **Female 5** | **157** | **202** | **143** | **153** | **227** | **227** | **173** | **173** | **216** | **218** |
|  | **Male 5** | **157** | **162** | **143** | **162** | **224** | **224** | **173** | **173** | **218** | **218** |
|  | **Seed 1** | 162 | 202 | 143 | 153 | 224 | 227 | 173 | 173 | 216 | 218 |
|  | **Seed 2** | 157 | 162 | 143 | 153 | 224 | 227 | 173 | 173 | 216 | 218 |
|  | **Seed 3** | 157 | 202 | 143 | 153 | 224 | 227 | 173 | 173 | 218 | 218 |
|  | **Seed 4** | 157 | 157 | 143 | 153 | 224 | 227 | 173 | 173 | 218 | 218 |
|  | **Seed 5** | 157 | 162 | 143 | 153 | 224 | 227 | 173 | 173 | 218 | 218 |
|  | **Seed 6** | 162 | 202 | 143 | 153 | 224 | 227 | 173 | 173 | 218 | 218 |
|  | **Seed 7** | 162 | 202 | 143 | 153 | 224 | 227 | 173 | 173 | 216 | 218 |
|  | **Seed 8** | 157 | 157 | 143 | 153 | 224 | 227 | 173 | 173 | 216 | 218 |
| **Couple 6** | **Female 6** | **171** | **201** | **143** | **151** | **221** | **224** | **173** | **173** | **218** | **218** |
|  | **Male 6** | **198** | **214** | **143** | **143** | **224** | **224** | **173** | **173** | **216** | **218** |
|  | **Seed 1** | 171 | 198 | 143 | 143 | 224 | 224 | 173 | 173 | 218 | 218 |
|  | **Seed 2** | 171 | 214 | 143 | 143 | 224 | 224 | 173 | 173 | 218 | 218 |
|  | **Seed 3** | 171 | 214 | 143 | 143 | 224 | 224 | 173 | 173 | 216 | 218 |
|  | **Seed 4** | 201 | 214 | 143 | 151 | 221 | 224 | 173 | 173 | 218 | 218 |
|  | **Seed 5** | 201 | 214 | 143 | 151 | 221 | 224 | 173 | 173 | 218 | 218 |
|  | **Seed 6** | 198 | 201 | 143 | 143 | 224 | 224 | 173 | 173 | 216 | 218 |
|  | **Seed 7** | 198 | 201 | 143 | 151 | 221 | 224 | 173 | 173 | 218 | 218 |
|  | **Seed 8** | 198 | 201 | 143 | 143 | 224 | 224 | 173 | 173 | 216 | 218 |
| **Couple 7** | **Female 7** | **162** | **198** | **143** | **151** | **224** | **224** | **173** | **173** | **216** | **218** |
|  | **Male 7** | **162** | **188** | **143** | **149** | **224** | **224** | **173** | **174** | **222** | **222** |
|  | **Seed 1** | 162 | 188 | 143 | 151 | 224 | 224 | 173 | 174 | 218 | 222 |
|  | **Seed 2** | 162 | 188 | 143 | 143 | 224 | 224 | 173 | 174 | 216 | 222 |
|  | **Seed 3** | 162 | 188 | 143 | 149 | 224 | 224 | 173 | 174 | 216 | 222 |
|  | **Seed 4** | 162 | 198 | 143 | 151 | 224 | 224 | 173 | 173 | 218 | 222 |
|  | **Seed 5** | 162 | 188 | 143 | 151 | 224 | 224 | 173 | 174 | 216 | 222 |
|  | **Seed 6** | 162 | 188 | 143 | 149 | 224 | 224 | 173 | 174 | 218 | 222 |
|  | **Seed 7** | 162 | 188 | 149 | 151 | 224 | 224 | 173 | 174 | 216 | 222 |
|  | **Seed 8** | 162 | 198 | 149 | 151 | 224 | 224 | 173 | 173 | 218 | 222 |
| **Couple 8** | **Female 8** | **160** | **200** | **151** | **155** | **224** | **224** | **173** | **173** | **218** | **222** |
|  | **Male 8** | **159** | **162** | **143** | **143** | **221** | **224** | **173** | **173** | **220** | **222** |
|  | **Seed 1** | 160 | 162 | 143 | 151 | 221 | 224 | 173 | 173 | 222 | 222 |
|  | **Seed 2** | 160 | 162 | 143 | 151 | 221 | 224 | 173 | 173 | 218 | 222 |
|  | **Seed 3** | 159 | 160 | 143 | 151 | 221 | 224 | 173 | 173 | 218 | 222 |
|  | **Seed 4** | 160 | 162 | 143 | 155 | 224 | 224 | 173 | 173 | 222 | 222 |
|  | **Seed 5** | 159 | 160 | 143 | 151 | 224 | 224 | 173 | 173 | 218 | 222 |
|  | **Seed 6** | 159 | 200 | 143 | 151 | 224 | 224 | 173 | 173 | 218 | 222 |
|  | **Seed 7** | 159 | 160 | 143 | 155 | 221 | 224 | 173 | 173 | 218 | 222 |
|  | **Seed 8** | 159 | 160 | 143 | 151 | 221 | 224 | 173 | 173 | 222 | 222 |
| **Couple 9** | **Female 9** | **172** | **178** | **141** | **143** | **224** | **227** | **173** | **173** | **218** | **220** |
|  | **Male 9** | **178** | **178** | **143** | **156** | **224** | **224** | **173** | **176** | **220** | **222** |
|  | **Seed 1** | 178 | 178 | 141 | 143 | 224 | 224 | 173 | 176 | 218 | 222 |
|  | **Seed 2** | 172 | 178 | 141 | 156 | 224 | 227 | 173 | 176 | 220 | 220 |
|  | **Seed 3** | 172 | 178 | 141 | 143 | 224 | 224 | 173 | 173 | 220 | 222 |
|  | **Seed 4** | 172 | 178 | 141 | 156 | 224 | 224 | 173 | 176 | 218 | 220 |
|  | **Seed 5** | 178 | 178 | 143 | 156 | 224 | 224 | 173 | 173 | 220 | 220 |
|  | **Seed 6** | 178 | 178 | 143 | 143 | 224 | 224 | 173 | 176 | 218 | 220 |
|  | **Seed 7** | 178 | 178 | 141 | 156 | 224 | 224 | 173 | 173 | 218 | 222 |
|  | **Seed 8** | 172 | 178 | 141 | 143 | 224 | 227 | 173 | 176 | 220 | 220 |
| **Couple 10** | **Female 10** | **149** | **178** | **143** | **145** | **221** | **224** | **173** | **173** | **218** | **222** |
|  | **Male 10** | **151** | **160** | **143** | **143** | **221** | **224** | **173** | **173** | **218** | **218** |
|  | **Seed 1** | 151 | 178 | 143 | 143 | 221 | 224 | 173 | 173 | 218 | 222 |
|  | **Seed 2** | 151 | 178 | 143 | 145 | 221 | 221 | 173 | 173 | 218 | 222 |
|  | **Seed 3** | 149 | 151 | 143 | 145 | 221 | 224 | 173 | 173 | 218 | 218 |
|  | **Seed 4** | 149 | 151 | 143 | 145 | 221 | 224 | 173 | 173 | 218 | 218 |
|  | **Seed 5** | 149 | 160 | 143 | 143 | 224 | 224 | 173 | 173 | 218 | 218 |
|  | **Seed 6** | 160 | 178 | 143 | 145 | 224 | 224 | 173 | 173 | 218 | 218 |
|  | **Seed 7** | 160 | 178 | 143 | 160 | 221 | 221 | 173 | 173 | 218 | 222 |
|  | **Seed 8** | 160 | 178 | 143 | 145 | 221 | 224 | 173 | 173 | 218 | 222 |
| **Couple 11** | **Female 11** | **178** | **178** | **143** | **143** | **224** | **224** | **173** | **173** | **218** | **222** |
|  | **Male 11** | **178** | **246** | **143** | **156** | **224** | **227** | **173** | **173** | **218** | **218** |
|  | **Seed 1** | 178 | 246 | 143 | 156 | 224 | 224 | 173 | 173 | 218 | 218 |
|  | **Seed 2** | 178 | 246 | 143 | 156 | 224 | 227 | 173 | 173 | 218 | 218 |
|  | **Seed 3** | 178 | 246 | 143 | 143 | 224 | 224 | 173 | 173 | 218 | 218 |
|  | **Seed 4** | 178 | 246 | 143 | 156 | 224 | 224 | 173 | 173 | 218 | 222 |
|  | **Seed 5** | 178 | 178 | 143 | 143 | 224 | 224 | 173 | 173 | 218 | 218 |
|  | **Seed 6** | 178 | 178 | 143 | 156 | 224 | 224 | 173 | 173 | 218 | 218 |
| **Couple 12** | **Female 12** | **200** | **202** | **145** | **151** | **221** | **224** | **173** | **173** | **220** | **222** |
|  | **Male 12** | **150** | **201** | **143** | **143** | **221** | **224** | **173** | **173** | **218** | **222** |
|  | **Seed 1** | 150 | 202 | 143 | 145 | 221 | 221 | 173 | 173 | 218 | 220 |
|  | **Seed 2** | 150 | 202 | 143 | 145 | 221 | 224 | 173 | 173 | 220 | 222 |
|  | **Seed 3** | 150 | 202 | 143 | 151 | 221 | 224 | 173 | 173 | 220 | 222 |
|  | **Seed 4** | 150 | 202 | 143 | 151 | 221 | 221 | 173 | 173 | 218 | 220 |
|  | **Seed 5** | 150 | 202 | 143 | 145 | 221 | 224 | 173 | 173 | 218 | 222 |
|  | **Seed 6** | 150 | 202 | 143 | 145 | 221 | 224 | 173 | 173 | 218 | 220 |
|  | **Seed 7** | 150 | 202 | 143 | 145 | 221 | 224 | 173 | 173 | 218 | 222 |
|  | **Seed 8** | 150 | 202 | 143 | 145 | 221 | 221 | 173 | 173 | 220 | 222 |
|  | **χ²_1,94_** | 0.800 |  | 1.138 |  | 1.543 |  | 3.112 |  | 1.543 |  |
|  | ***p*** | 0.370 |  | 0.286 |  | 0.214 |  | 0.078 |  | 0.214 |  |

***References***

Barbot, E., Dufaÿ, M. & De Cauwer, I. (2023) Sex-specific selection patterns in a dioecious insect-pollinated plant. *Evolution,* **77,** 1578–1590.

Barbot, E., Dufaÿ, M., Godé, C. & De Cauwer, I. (2025) Investigating the effects of diurnal and nocturnal pollinators on male and female reproductive success and on floral trait selection in *Silene dioica Peer Community Journal* **5**.

Barbot, E., Dufaÿ, M., Tonnabel, J., Godé, C. & De Cauwer, I. (2022) On the function of flower number: disentangling fertility from pollinator-mediated selection. *Proceedings of the Royal Society B: Biological Sciences,* **289,** 20221987.

Baskin, C. C. & Baskin, J. M. (2019) Martin's peripheral embryo – unique but not a phylogenetic ‘orphan’ at the base of his family tree: a tribute to the insight of a pioneer seed biologist. *Seed Science Research,* **29,** 155-166.

Broyles, S. B. & Wyatt, R. (1990) Paternity analysis in a natural population of *Asclepias exaltata*: multiple paternity, functional gender, and the "Pollen-Donation Hypothesis". *Evolution,* **44,** 1454-1468.

Kartal, C. & Tekin, M. (2021) Embryological studies on *Silene muradica* Schischk. (Caryophyllaceae) – A gynodioecious species from Turkey *Acta Biologica Cracoviensia,* **63,** 21-29.

Martin, A. C. (1946) The comparative internal morphology of seeds. *The American Midland Naturalist,* **36,** 513-660.

Meagher, T. R. (1986) Analysis of paternity within a natural population of *Chamaelirium luteum*. 1. Identification of most-likely male parents. *American Naturalist,* **128,** 199-215.

Mohana Rao, P. R., L., G. J. & and Duret, S. (1988) An ultrastructural study of perisperm and endosperm in *Silene alba* Miller E.H.L. Krause. *Bulletin de la Société Botanique de France. Lettres Botaniques,* **135,** 123-130.

**Method S2. Calculation of dissimilarity coefficients.**

In this study, we quantified sire profile similarity for each pair of fruits by calculating beta diversity indices following Legendre (2014), using both a qualitative and a quantitative version of the incidence matrix, with fruits arranged in rows and pollen donors arranged in columns (see below for examples).

*Qualitative incidence matrix*

|  | **Male 1** | **Male 2** | **Male 3** | **Male 4** |
| --- | --- | --- | --- | --- |
| **Fruit A** | 1 | 1 | 1 | 0 |
| **Fruit B** | 1 | 0 | 0 | 1 |

From the qualitative matrix, we calculated a as the number of sires detected in both fruits, b as the number of sires detected in fruit A but not in fruit B and c as the number of sires detected in fruit B but not in fruit A. In the example above, a = 1, b = 2 and c = 1.

We then computed the Sørensen index D_S_ = (b+c)/(2a+b+c).

This index can be further partitionned into two components:

-the proportion of dissimilarity due to remplacement of a given sire by another one, Repl_S_ =|b-c|/(2a+b+c). In the example above, Repl_S_ = 0.40.

-the proportion of dissimilarity due to differences in paternal diversity between fruits A and B, Rich_S_ = 2×min(b,c)/(2a+b+c). In the example above, Rich_S_ = 0.20.

*Quantitative incidence matrix*

|  | **Male 1** | **Male 2** | **Male 3** | **Male 4** |
| --- | --- | --- | --- | --- |
| **Fruit A** | 8 | 4 | 2 | 0 |
| **Fruit B** | 10 | 0 | 0 | 4 |

From the quantitative matrix, we calculated A as the sum of minimum abundances (*i.e.*, we retained the lowest number of seeds assigned to each sire between fruits A and B, and summed these minima across all sires), B as the sum of abundance differences in fruit A (*i.e.*, we calculated the number of seeds assigned to each sire minus the minimum, and summed these differences accross all sires) and C as the sum of abundance differences in fruit B (*i.e.*, calculated as explained for fruit A). In the example above, A = 8, B = 6 and C = 6.

We then computed the quantitative version of the Sørensen index, that is, the percentage difference, D_%diff_ = (B+C)/(2A+B+C).

This index can be further partitionned into two components:

-the proportion of dissimilarity due to replacement of seeds assigned to a given sire by seeds assigned to another one, Repl_%diff_ = 2×min(B,C)/(2A+B+C). In the example above, Repl_%diff_ = 0.43.

-the proportion of dissimilarity due differences in the total number of seeds considered for fruits A and B, AbDiff_%diff_ = |B-C|/(2A+B+C). In the example above as in our dataset, the total number of seeds considered was standardized across fruits using a bootstrapping procedure, so that AbDiff_%diff_ = 0.

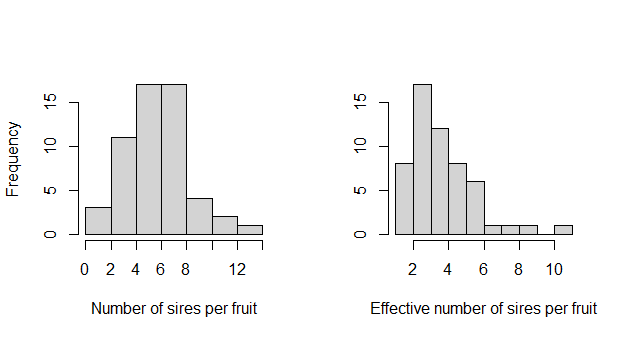

**Figure S1.** Distribution of the number of sires per fruit (lef panel) and effective number of sires (right panel) in the 55 S. dioica fruits sampled in this study.

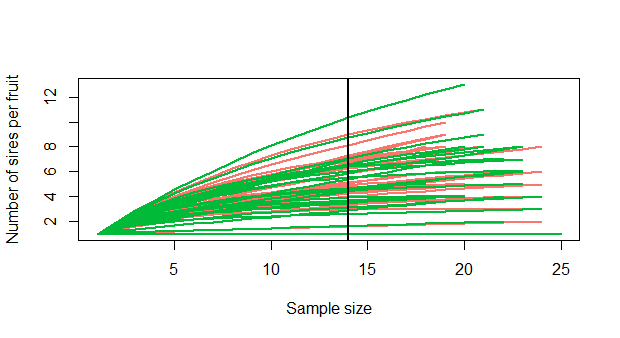

**Figure S2.** Rarefaction curves for the 55 fruits sampled in this study. Fruits sampled on manipulated plants are shown in cyan, and fruits sampled on control plants in magenta. The vertical line corresponds to the sample size of 14 used in this study.
